# Lifetime of a structure evolving by cluster aggregation and particle loss, and application to postsynaptic scaffold domains

**DOI:** 10.1101/2019.12.27.889196

**Authors:** Vincent Hakim, Jonas Ranft

## Abstract

The dynamics of several mesoscopic biological structures depend on the interplay of growth through the incorporation of components of different sizes laterally diffusing along the cell membrane, and loss by component turnover. In particular, a model of such an out-of-equilibrium dynamics has recently been proposed for postsynaptic scaffold domains which are key structures of neuronal synapses. It is of interest to estimate the lifetime of these mesoscopic structures, especially in the context of synapses where this time is related to memory retention. The lifetime of a structure can be very long as compared to the turnover time of its components and it can be difficult to estimate it by direct numerical simulations. Here, in the context of the model proposed for postsynaptic scaffold domains, we approximate the aggregation-turnover dynamics by a shot-noise process. This enables us to analytically compute the quasi-stationary distribution describing the sizes of the surviving structures as well as their characteristic lifetime. We show that our analytical estimate agrees with numerical simulations of a full spatial model, in a regime of parameters where a direct assessment is computationally feasible. We then use our approach to estimate the lifetime of mesoscopic structures in parameter regimes where computer simulations would be prohibitively long. For gephyrin, the scaffolding protein specific to inhibitory synapses, we estimate a lifetime longer than several months for a scaffold domain when the single gephyrin protein turnover time is about half an hour, as experimentally measured. While our focus is on postsynaptic domains, our formalism and techniques should be applicable to other biological structures that are also formed by a balance of condensation and turnover.

## I. INTRODUCTION

Synapses play a central role in learning and memory. The determination of their components, of their structure and of their dynamics has been the focus of numerous investigations. On the postsynaptic neurotransmitter-receiving side of the synapse, the synaptic proteins forming the postsynaptic density (PSD) are structurally organized by scaffolding proteins. Biophysical experiments have demonstrated that synapses are very dynamic structures [1]. Neurotransmitter receptors are transmembrane proteins that have been imaged to go in and out of synapses by single particle tracking techniques. The scaffolding proteins themselves have been shown to turn over at synapses on the timescale of an hour. As anticipated by Francis Crick 35 years ago [2], this raises the fundamental and still unanswered question of how long-term memory persists for years.

Motivated by this puzzle and the wealth of quantitative biophysical data, we proposed in a previous work [3] a simple biophysical model for the formation of postsynaptic scaffold domains, taking inhibitory synapses as a starting point. For inhibitory synapses, the scaffolding protein gephyrin can form large oligomers through homophilic binding at its two ends. (Gephyrin proteins naturally occur as trimers that can oligomerize via the three remaining dimeric binding sites [4].) Large assemblies of the gephyrin scaffolding protein are located just below the postsynaptic membrane [4] and serve as structural foundation to the PSD. As gephyrin can bind transmembrane inhibitory neurotransmitter receptors such as GABA and glycine receptors on their cytoplasmic side, receptor concentration at inhibitory synapses is increased 20-50 fold as compared to the concentration of receptors laterally diffusing in the extrasynaptic membrane [5, 6].

Available data [1, 7] suggest that both receptors and gephyrin scaffolding proteins are transported to the extrasynaptic cell membrane. Data also suggests [5] that gephyrin scaffolding proteins, that are not membrane proteins, diffuse laterally along the membrane, carried along by their binding to the cytoplasmic domain of glycine receptors. Relying on this experimental evidence, we proposed that postsynaptic scaffold domains are formed and maintained by the continuous “aggregation”, i.e. encounter and oligomerization, of scaffolding proteins that laterally diffuse as receptor-scaffold complexes, counteracted by the desorption of scaffolding proteins into the cytoplasm [3]. A sketch of the proposed aggregation-removal dynamics is shown in Fig. 1A. The possible aggregation of diffusing complexes outside of synapses led us to predict that scaffold domains with a continuous range of sizes should be observed extrasynaptically, as illustrated in the simulation snapshot shown in Fig. 1B. More quantitatively, numerical simulations produce a distribution of concentrations *c_n_* of clusters of *n* particles (Fig. 1C), which is consistent with reported data on gephyrin clusters [3, 8].

**FIG. 1.**
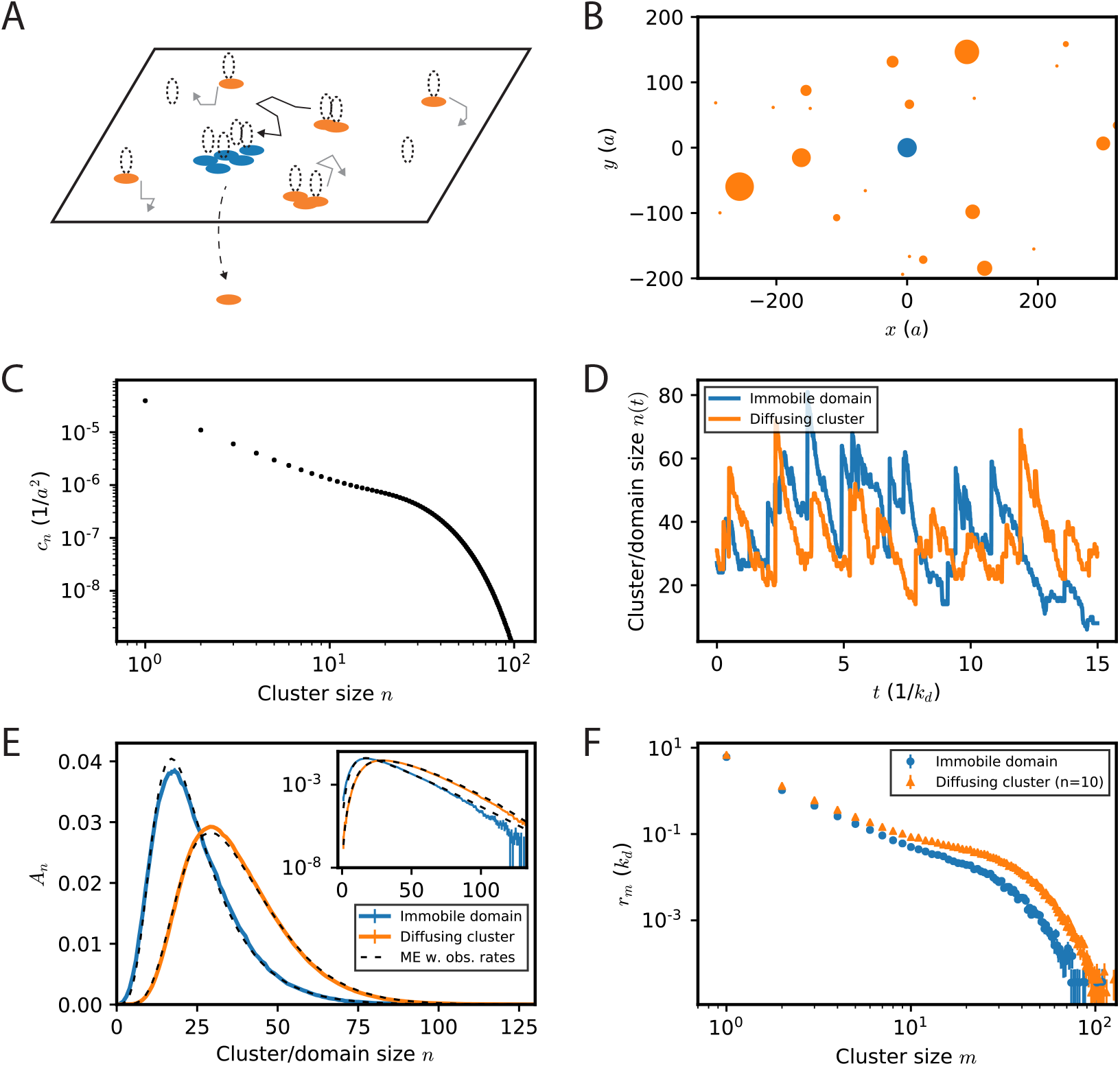
Dynamics of diffusing clusters and of an immobile domain of fluctuating size. (A) Schematic description of the dynamics of postsynaptic scaffold clusters and of an immobile scaffold domain. Clusters of scaffolding proteins (orange) diffuse laterally under the membrane, complexed with neurotransmitter receptors (dashed ellipses). Upon encounter, the diffusing clusters can aggregate and grow. They can also encounter an immobile domain (blue) of fluctuating size meant to represent the postsynaptic scaffold domain of a synapse. The growth of the diffusing clusters and of the immobile domain is counter-balanced by the turnover of scaffolding proteins into the cytoplasm. (B) Snapshot of a simulation of diffusing and aggregating particles (orange) subject to turnover [3], with an immobile domain of fluctuating size (blue). For illustrative purposes the cluster and domain sizes are scaled by a factor of five with respect to their actual sizes. The diameter *a* of a single particle is taken as length unit. Taking *a* as the distance between two neighboring vertices in a hexagonal lattice of gephyrin trimers gives *a* 20 nm. (C) Stationary concentrations *c_n_* of diffusing clusters of size *n* (*n* being the number of constituent particles) resulting from the simulated aggregation-turnover dynamics. (D) Example of the evolution over time of the sizes of the immobile domain (blue) and of a freely diffusing (orange) cluster that grow by stochastic encounters with other clusters and shrink by particle desorption. Fusion with impinging clusters of different sizes lead to instantaneous size increases, while Poissonian particle loss produces slower size decreases. (E) The probability distributions *A_n_* of the sizes *n* of an immobile domain (solid blue line) and of long-lived diffusing clusters (solid orange line) appear stationary after a few turnover times 1*/k_d_*. The panel also shows the corresponding quasi-stationary distributions obtained from the master equations (ME) for immobile domains (Eq. (42), dashed blue line) and diffusing clusters (Eq. (51), dashed orange line), corresponding to the shot-noise model proposed in the present work, where the size dynamics are based on the rates of impingement shown in F. (F) Encounter rates *r_m_* of the immobile domain with diffusing clusters of size *m* as measured in the simulation (blue dots). The encounter rates with diffusing clusters of different sizes *m* are also shown (orange) for a freely diffusing cluster followed over time. In this case, the encounter rates as well as the diffusion constant (Eq. (45)) of the followed cluster depends on its size *n* (here *n* = 10, *σ* = 0.5).

Focusing on a single synaptic domain, a steady state is attained when, on average, particle loss due to turnover is balanced by the fusion with extrasynaptic protein aggregates of all sizes. In numerical simulations, the domain size appears constant on average, but its size fluctuates due to both stochastic particle loss and stochastic fusion with surrounding diffusing clusters, as illustrated in Fig. 1D. After a few single particle turnover times, the probability of the different possible sizes appears well-described by a stationary probability distribution (Fig. 1E). Eventually however, rare fluctuations should lead to a decrease of the domain size followed by its disappearance. The distribution of fluctuating domain sizes shown in Fig. 1E should thus be more accurately described as a quasi-stationary distribution, namely the distribution of domain sizes conditioned on the survival of the domains [9].

Although the dynamics of an immobile domain is our main interest, it is also possible to consider the dynamics of a diffusing cluster of particles that is followed over time [3]. The process is qualitatively very similar to that for an immobile domain, as shown in Fig. 1D. The quasistationary distributions of sizes in the two cases are also similar (Fig. 1E) and very different from the distribution of cluster sizes at a given time (Fig. 1C). Please note that here and in the following, we call “domains” aggregates that do not diffuse and “clusters” aggregates that diffuse laterally along the membrane (bound to receptors that diffuse in the membrane, in the case of scaffolding proteins that are our main focus here).

In the present study, our aim is to obtain the quasistationary size distribution and to estimate the lifetime of such a domain of mesoscopic size evolving by stochastic aggregation with clusters of a range of sizes *m* and Poissonian single particle loss. In section II A, we introduce and analyze a simplified model for the fluctuating domain dynamics in which we simply retain the rate *r_m_* (Fig. 1F) of clusters of different sizes aggregating with the immobile fluctuating domain, reducing the impinging dynamics to a shot-noise process [10]. This amounts to neglecting all correlations between the impingement times of different clusters. In this formulation, obtaining the time of first passage to zero of the domain size, is conceptually similar to determining the time of extinction of a population in a birth-death rate process. We employ master equation techniques similar to the ones used in that context [11] to obtain the quasi-stationary size distribution of the fluctuating domains as well as their lifetime. We first consider the case when impinging clusters are monodisperse in section II B and then generalize our analysis to polydisperse clusters in section II C. We then show in section III A that the proposed simplified model provides a good approximation to the quasi-stationary size distribution and lifetime of domains measured in numerical simulations of diffusing clusters. This requires to choose kinetic parameters for which the cluster survival is long compared to the single molecule turnover time but short enough to be observed and quantified in numerical simulations. We also briefly discuss in this setting, in section III B, the more complicated case of a diffusing cluster that is followed over time. Finally, in section III C, biological parameters appropriate for gephyrin scaffold domains are used to provide a theoretical estimation of their lifetime, based on our previously proposed description of their dynamics [3].

Our focus in this paper is on postsynaptic domains but other membrane-less condensed structures have recently sparked much interest. Some of them have been viewed as condensed structures in thermodynamic equilibrium [12, 13]. Others, such as lipid rafts [14, 15] and E-cadherin clusters [16] have been proposed to be formed and maintained as non-equilibrium steady states by condensation and recycling of components, somewhat analogously to the model for postsynaptic scaffold domains proposed in ref. [3]. These latter schemes consume energy but provide, for instance, means to tune the size of the condensed structure independently of the size of the cellular compartment which contains it. The methods used in the present paper should be generalizable to these other cases.

## II. SIMPLIFIED DYNAMICS OF AN IMMOBILE DOMAIN

We consider the evolution of an immobile domain, the size of which grows by stochastic aggregation of clusters of particles and diminishes due to single particle loss. We suppose that clusters of size *m* impinge stochastically on the domain and fuse with it at a rate *r_m_* in a Poissonian fashion. Particle loss from the domain is also supposed to be stochastic and Poissonian at a rate *k_d_*. When cluster aggregation dominates over particle loss for small domain sizes, domains typically fluctuate around a large mean size ⟨n⟩ ⨠ 1, for a long time as compared to 1*/k_d_*, before eventually losing all their particles and disappearing. Our aim in the present section is to analyze this process, inspired by the process illustrated in Fig. 1 A, and in particular to precisely evaluate the domain survival time.

### A. Large domains and shot-noise processes

In order to provide some feeling for the considered shot-noise process, we assume in this first subsection that the typical size of the domain is large and neglect the discreteness of particle loss [3]. The size discreteness is also lost in this description which makes us denote size by *s* instead of *n*.

The evolution of the domain size *s* can thus approximately be described as

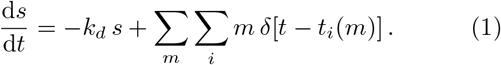

The first term on the right-hand side (r.h.s.) of Eq. (1) describes the domain size reduction due to single particle loss at a rate *k_d_*. We assume that all particles forming the domain have the same probability to leave it, which gives a total loss proportional to the domain size *s*. The second term on the r.h.s. describes the sudden domain size increases resulting from its encounters with clusters of size *m* at times {*t_i_*(*m*), *i* = …, −1, 0, 1, …}. The times *t_i_*(*m*) have been indexed by the size of the impinging clusters since we suppose that the corresponding events occur in a Poissonian fashion at rates *r_m_* that only depend on the impinging cluster size *m*. When particle loss is described in a continuous way as in Eq. (1), the domain size *s* is always strictly positive and its lifetime is infinite. This continuous approximation nonetheless accurately describes the typical size of long-lived clusters and regular fluctuations around it. We thus first consider it to obtain some basic characteristics of the process described by Eq. (1) which belongs to a class of well-studied shot-noise evolutions [17].

With Eq. (1), the mean domain size ⟨*s*⟩ is simply obtained as a balance between the particle loss flux, −*k_d_*⟨*s*⟩, and the flux ∑_*m*_ *r_m_m* of clusters impinging on the domain,

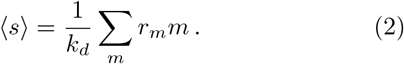

Other moments of the domain size can also easily be computed for the linear Eq. (1). First, it is convenient to express *s*(*t*) in terms of a specific realization of the stochastic times {*t_i_*(*m*)},

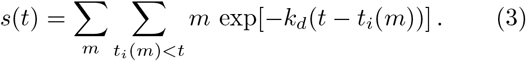

The moments can then be obtained by averaging over the stochastic times *t_i_(m)*. One obtains for the first few ones

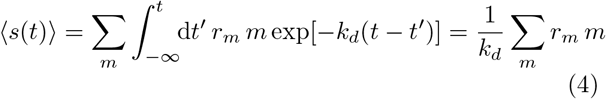

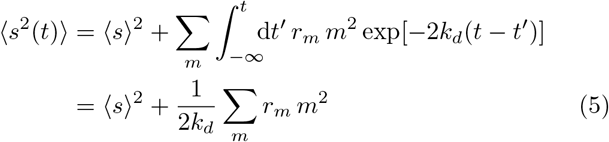

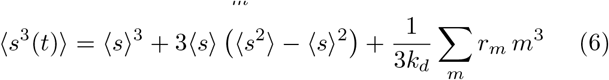

Since the average size ⟨s(t)⟩ is independent of time, it is simply denoted it by ⟨s⟩ in Eqs. (5,6). Correlation functions of low order can be computed in the same way,

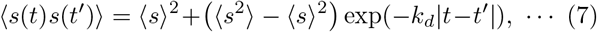

It is immediately apparent on the time series of the domain size shown in Fig. 1D that the process dynamics with its sharp upwards jumps and slower decreases, is not invariant under time reversal. This can be explicitly demonstrated, in the present framework, by showing that the third-order correlation function [18] ⟨*s(t)s(t′)^2^ − s(t)^2^s(t′)*⟩ does not vanish in the stationary state. Indeed, the explicit computation [3] gives

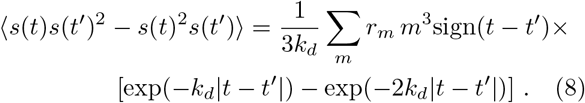

This absence of time-reversal invariance of correlation functions in the steady state shows that Eq. (1) does not describe the dynamics of a process at thermodynamic equilibrium [19] and that energy should be consumed to sustain it.

Higher correlation functions can also be computed. The computation of the *j*-th correlation function ⟨*s*(*t*_1_) … *s*(*t_j_*)⟩ involves sums over *j* stochastic times coming from the replacement of each *s*(*t_i_*) by its expression (3). The only term at order *j* that cannot be expressed with the help of of lower order correlation functions is obtained when the *j* stochastic times are identical in the sums. In other terms, the cumulants *c_j_* associated to the moments (*s^j^*) have the simple expression

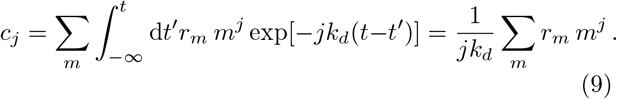

Thus the cumulant generating function G(*μ*) reads

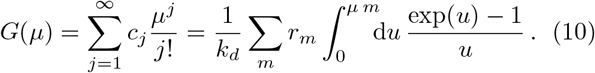

The moments are given by the corresponding generating function [17],

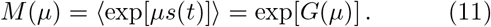

Another way [17] to analyze Eq. (1) and obtain Eq. (11) is to write the corresponding evolution equation for the probability *P* (*s, t*) of observing a cluster of size *s*,

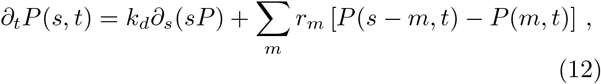

with *P* (*s, t*) = 0 for *s* < 0, where we have explicitly noted the convention used in Eq. (12) that *P* (*s, t*) vanishes for negative sizes. By taking the Laplace transform of *P* (*s*), Eq. (12) gives back Eq. (11), as we show below for the discrete case. Although we do not make use of it, we also note that the stationary version of Eq. (12) allows one [17] to explicitly compute the expression of *P* (*s*) in the successive size intervals, 0 *< s ≤ m, m < s* 2*m*, ….

When the fluctuations of *s*(*t*) are small compared to its average (i.e. *c*_2_ = ⟨*s*^2^(*t*)⟩ − ⟨*s*(*t*)⟩^2^) ≪ ⟨*s*(*t*)⟩^2^), it is clear that the domain size *s* will rarely become small and that its lifetime will be long as compared to 1*/k_d_*. In order to obtain a precise estimate, we turn in the following sections to the analysis of shot-noise process with discrete particle loss.

### B. Aggregation of monodisperse clusters

We begin by considering the simplest case when the impinging clusters have a single size *m*. Similarly to Eq. (12), master equations can easily be written for the probability *P_n_*(*t*) that the evolving domain has the size *n* at time *t*,

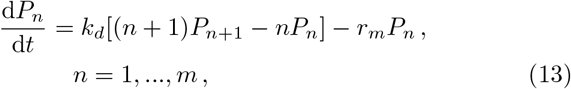

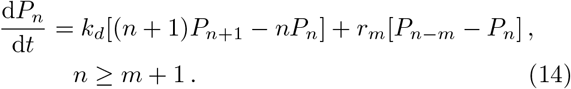

The first two terms on the r.h.s. of Eqs. (13,14) describe the domain size change due to particle loss. Namely particle loss at rate *k_d_* can produce a domain of size *n* from a domain of size *n* + 1 or change the size of an existing domain of size *n*. The other terms on the r.h.s. of Eq. (13,14) depict size change due to aggregation of a cluster of size *m* with rate *r_m_*: this can produce a cluster of size *n* from a cluster of size *n − m* if *n ≥ m* + 1, and change the size of an existing cluster. Fig. 2A shows an example trajectory with *m* = 10, *r_m_/k_d_* = 2 obtained from the simulation of the stochastic process described by Eq. (13,14).

**FIG. 2.**
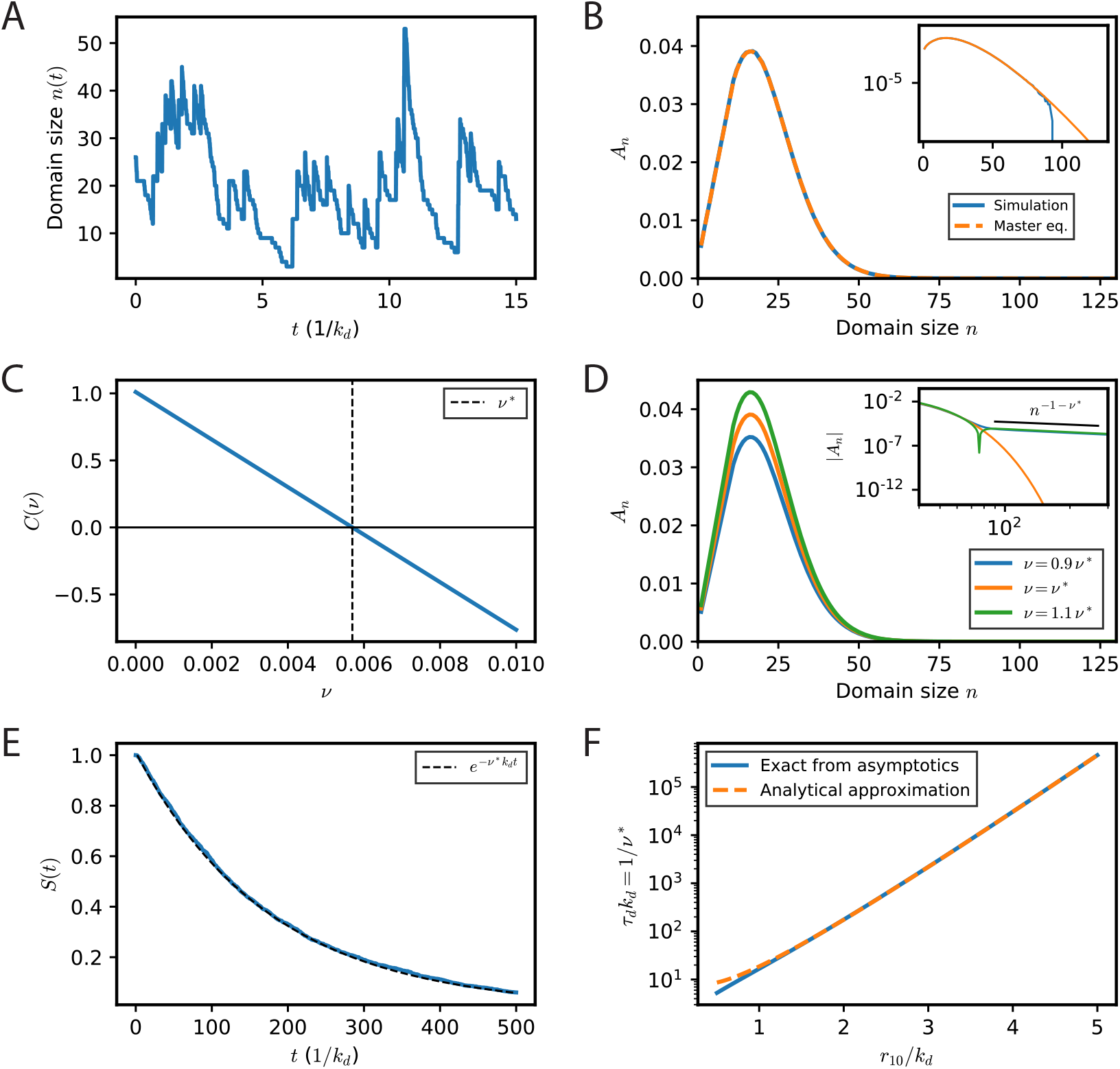
Shot-noise model for the stochastic size dynamics of an immobile domain in the case of mono-disperse impinging clusters. (A) Example of size evolution for a single domain described by the simplified shot-noise model. The randomly impinging clusters contain 10 particles, with an encounter rate relative to particle loss *r/k_d_* = 2 (the simulation shown was performed with *r* = 2, *k_d_* = 1). (B) The distribution of sizes explored by clusters after a transient time of 5*/k_d_* is shown. The distribution (solid blue line) represents the average over many (*n* = 2000) realizations of the stochastic shot-noise process for a duration of *T* = 500*/k_d_* each. The theoretical quasi-stationary distribution *A_n_* is also shown (dashed orange line). Its determination is explained in panels C & D. (C) The solution *A_n_* to the master equations (13,14) that govern the domain size distribution depends on the parameter *ν*. The slow *n*^−(1+*ν*)^ large-*n* asymptotics of the *A_n_* is proportional to *C*(*ν*) (Eq (26)) shown here. The sought-for solution of the master equations is obtained for *ν* = *ν*\*, when *C*(*ν*\*) = 0. (D) Three solutions of the master equations with different values of *ν* are shown, *ν* = *ν*\* = 5.69 · 10^−3^ (orange), *ν*_−_ = 0.9*ν*\* (blue), *ν*_+_ = 1.1*ν*\* (green). Inset: The distributions corresponding to *ν*_−_ and *ν*_+_ have a slow decay for large *n* which agrees with the predicted scaling *n*^−(1+*ν*)^. The distribution for *ν* = *ν*\* decays much faster. All distributions are normalized to one. The contributions of the slow positive tail for *ν*_−_ and slow negative tail for *ν*+ explain their apparent differences in normalization. (E) The number of surviving clusters (solid blue line) in the simulations agrees well with the predicted exponential decay exp(−*ν*\**k_d_t*) (dashed black curve). (F) The characteristic lifetime *τ_d_* of the clusters increases exponentially with the rate of supply *r*_10_. The theoretical prediction of the lifetime with the numerically determined *ν*\* (blue line) is well approximated by the analytical expression (Eq. (20)) for *r*_10_*/k_d_* ≳ 1 (orange dashed line).

When *r_m_ ≫ k_d_*, the balance between cluster aggregation and particle loss produces domains fluctuating for a long time around the large size *mr_m_/k_d_*. Eventually, however, the domain comprises only one particle and has a finite probability of disappearing. This separation of timescales renders very relevant the mathematical notion of quasi-stationary distribution [9], that is a probability distribution that is stationary when the domain is conditioned not to disappear. For *r_m_ ≫ k_d_*, the *P_n_* relax on a fast timescale of order 1*/k_d_* to a quasi-stationary distribution, an example of which is shown in Fig. 2B. The probability to remain in this quasi-stationary distribution then slowly decreases. We compute below the smallest eigenvalue of the system (13,14) which characterizes this slow decrease and provides the (inverse) timescale of domain disappearance. Similar estimates together with rigorous proofs have been provided before for birth-and-death processes in the context of population dynamics [11]. We describe first another way to estimate the small flux of disappearing domains. It consists in turning the process into a stationary one by fictitiously recreating a domain after its disappearance. This is similar in spirit to Kramers’s classic method [20] of obtaining the time of thermal escape out of a finite potential well from the steady state particle flux produced by re-injecting particles into the well after their escape.

#### 1. A stationary process with domain recreation

In order to transform Eqs. (13,14) into a stationary process, we monitor the creation of a domain of vanishing size by introducing the probability *P*_0_ that a domain contains 0 particles. We also allow the creation of a domain of size *m* from a domain with zero particles. Namely, we supplement (13,14) with

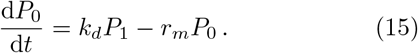

At the same time, Eq. (14) is supposed to be valid starting from *n* = *m* (instead of *n* = *m* + 1) with *P*_0_ now well-defined. In other words, a domain that has disappeared is reinjected at a size *m* with a rate *r_m_*. This prescription turns the domain dynamics into a process that has a well-defined steady state.

Setting the time derivatives to zero, Eq. (15) determines *P*_1_ from *P*_0_ and by recursion all the *P_n_* with the help of Eqs. (13,14). One way to solve these equations is to introduce the generating function for the *P_n_*,

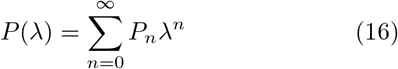

which obeys the differential equation

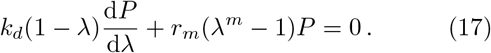

One can note that the definitions of *P* (*λ*) coincide with that of the previously defined *M* (*μ*) (Eq. (11)) with *λ* = exp(*μ*). The solution of Eq. (17) is readily obtained as

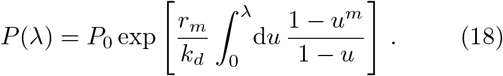

Requiring that the sum of the *P_n_* equals 1 determines *P*_0_ as well as the mean probability flux *J_d_* for a single cluster to reach a size 0,

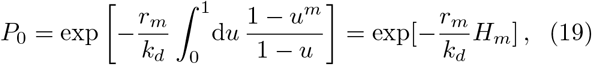

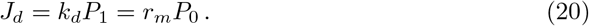

where we have used that the integral in Eq. (19) is the integral representation of the harmonic number *H_m_*,

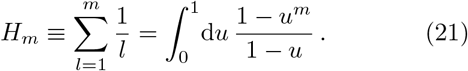

Expansion in powers of *λ* of *P* (*λ*) (Eq. (18,19)) provides explicit expressions for the *P_n_*. The large-*n* asymptotics of the *P_n_* can also be determined from the large-*λ* behavior of *P* (*λ*) (see Appendix A). The inverse of *J_d_* provides the sought-after estimation of the mean domain disappearance time *τ_d_*.

In the limit of large *m*, one can check that the expression (18) gives back the previously found expression for *M* (*μ*) when applied to *P* [exp(*μ*)]. In the same limit, the mean domain disappearance time *τ_d_* can be simply expressed.

For large *m*, the harmonic number *H_m_* can be approximated as *H_m_* ≃ ln *m* + *γ*, where *γ* ≃ 0.577 is Euler’s constant. The mean domain disappearance time thus reads in the large *m* limit,

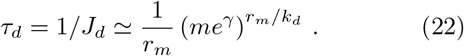

#### 2. Computation of the smallest relaxation eigenvalue

Another estimate of the domain disappearance time is provided by the maximal eigenvalue [11] associated to the linear evolution (13,14). Namely, we search the smallest positive *ν* and allied eigenvector *A_n_*, such that *P_n_* = *A_n_* exp(−*νk_d_t*) is an exact solution of the dynamics. Substitution of this ansatz for the *P_n_* in Eq. (13,14) provides explicit recursion formulas for the *A_n_* for a given *ν*,

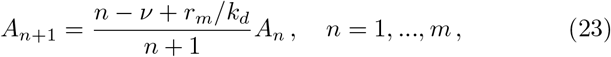

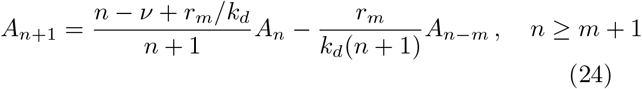

These recursion relations determine all the *A_n_* as a function of *A*_1_ for a given *ν*. The admissible values of *ν* are themselves determined by considering the asymptotic behavior of the *A_n_* for *n* large. Assuming that *A_n_* varies slowly with *n*, Eq. (24) gives (for *n* ≫ 1),

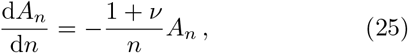

that is,

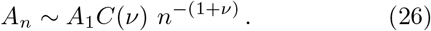

The numerical iteration of the recursion relations (23,24) indeed produces the slow asymptotic decay (26) for general values of *ν*. The obtained prefactor *C*(*ν*) is displayed in Fig. 2C. The slow asymptotic decay (26) does not correspond to the solution we seek. The function *C*(*ν*) is found to be to be negative for *ν > ν*\* (Fig. 2C). Therefore, all the values of *ν* such that *ν > ν*\* give negative probabilities and can be immediately excluded. The other values, *ν < ν*\*, with slow decay can also be excluded since for *n* large, the incoming flux of clusters of size *n* is dominated by the decaying larger clusters and not the growing smaller clusters (i.e. *k_d_*(*n* + 1)*P_n_*_+1_ *> r_m_P_n_*_−*m*_). This shows that these solutions correspond to initial cluster distributions with long tails of large clusters and not to the distribution we are interested in which is created by the growth of small clusters. Additionally, when *ν*\* < 1, as we will obtain, the slow asymptotic decay Eq. (26) gives distributions with diverging mean cluster sizes when *ν* < *ν*\*. This divergence is not possible when large clusters are produced from aggregation of smaller clusters during a finite time before disappearing (and it is of course different from the mean size ≃ *mr_m_/k_d_* that we expect).

The sought-for maximal eigenvalue corresponds to *ν* = *ν*\* for which *C*(*ν*\*) = 0. In this case, the *A_n_* are positive for all *n* and have the much faster decay with ln(*A_n_*) ~ − [*n* ln(*n/a*) − *n*]*/m* (see Appendix A). Examples of the obtained *A_n_* for three different values of *ν* are shown in Fig. 2D, as well as the much faster asymptotic decay of the *A_n_* for *ν* = *ν*\*.

An explicit expression for *ν*\* can be obtained along lines similar to those of the previous section II B 1. It is convenient to introduce the generating function for the *A_n_*,

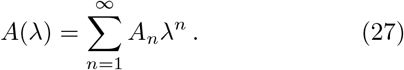

The recursion relations (23,24) translate into the following differential equation for *A*(*λ*),

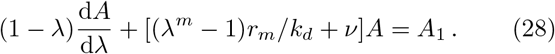

The solution of the linear Eq. (28) is easily obtained and reads

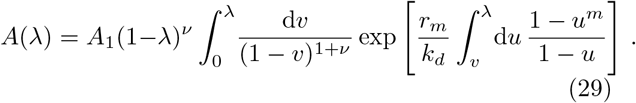

The singularity of *A*(*λ*) at *λ* = 1 determines the asymptotic behavior of the *A_n_*. Developing the above integral around *λ* = 1, one finds, as detailed in Appendix B,

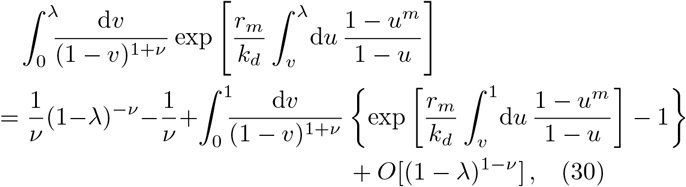

At *λ* = 1, *A*(*λ*) therefore behaves like

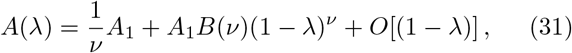

with

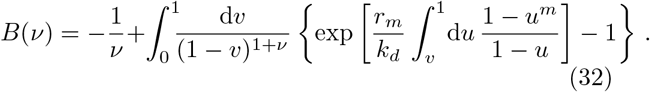

It is useful to remember that the coefficients *a_n_* of *λ^n^* in the expansion of (1 − *λ*)^*σ*^ are

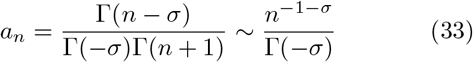

for large *n*, where Γ denotes the Gamma function [21]. Comparing Eq. (31) and (33) gives back the asymptotic behavior (26) of the *A_n_* with the explicit expression of *C*(*ν*) = *B*(*ν*)*/*(*ν*Γ(−*ν*)), or

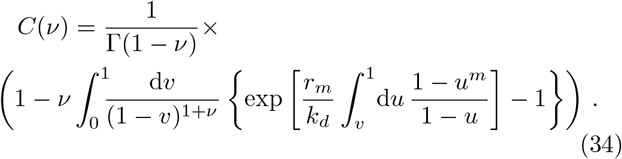

The function *C*(*ν*) (Fig. 2D) quickly decays with *ν* from *C*(0) = 1, vanishes at *ν* = *ν*\* and then becomes negative, reflecting the unphysical appearance of negative probabilities for *ν > ν*\*, as mentioned above. The starting value *C*(0) = 1 is clearly seen from the explicit expression (34). Its physical origin is simply cluster number conservation: For ν = 0, a stationary probability distribution is obtained by the injection of clusters of infinite size with the flux lim_*n*→+∞_(*k_d_ n A_n_*) = *k_d_C*(0)*A*_1_, which should be equal to the disappearance rate *k_d_A*_1_ of clusters of size 1.

In general, the equation *C*(*ν*\*) = 0 is an implicit equation for *ν*\*. However, in the limit where *r_m_m/k_d_* is large, the integral term in (34) is dominated by its lower bound. In this limit, one obtains the explicit asymptotic formula (see Appendix B)

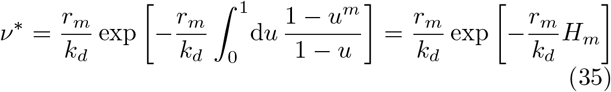

with *H_m_* given by Eq. (21). Eq. (35) gives the characteristic time *τ_d_* = 1*/*(*k_d_ν*\*) for the mean domain disappearance time. It is identical to our previous estimate (22). This analytical expression captures well the decay of the fraction of surviving clusters observed in simulations of the underlying shot-noise process, as shown in Fig. 2E & F (see Appendix C for details of the implementation).

### C. Aggregation of polydisperse clusters

We now extend the preceding calculation to the case of multiple aggregating clusters of different sizes *m*, with Poissonian encounter rates *r_m_*. These rates can be expressed in term of the concentrations *c_m_* of diffusing clusters of size *m* and their diffusion constant *D_m_*, as detailed in the next section. In the polydisperse case, the probabilities *P_n_* of the different sizes *n* of the fluctuating immobile domain evolve according to the master equations,

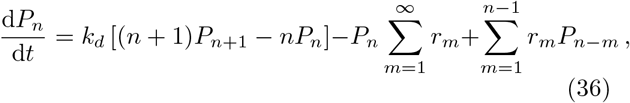

*n*≥1, where we have adopted the convention that *P_n_* = 0 for *n* ≤ 0, in the last term on the r.h.s.

We again search for the smallest positive eigenvalue *ν* and eigenvector *A_n_* such that *P_n_*(*t*) = *A_n_* exp[−*νk_d_ t*] is a solution of Eq. (36). Along the lines of the previous section, we introduce the generating function 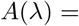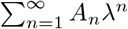, which now obeys the following differential equation:

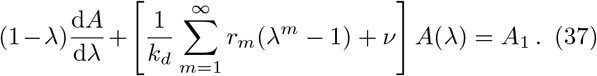

From this equation we can already deduce the average domain size ⟨⟨*n*⟩⟩ = ∑_*n*_ nP_n_(*t*)/ ∑_n_ P_n_(*t*) of surviving clusters, which is constant in the quasi-stationary regime and given by ⟨⟨*n*⟩⟩ = *A*′(1)/*A*(1) when expressed in terms of the generating function. Differentiating Eq. (37) with respect to *λ*, and setting *λ* = 1, one obtains −(1 − *ν*)*A*′(1) + ∑_*m*_(*r_m_/k_d_*)*m A*(1) = 0. For *ν* ≪ 1, we ⟨⟨*n*⟩⟩ ≃ ∑_*m*_ *r_m_m/k_d_*, consistent with the result obtained for the simplified description of section II.

The solution of Eq. (37) is again found relatively straightforwardly, and we obtain

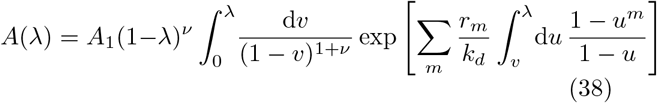

Following the arguments of the preceding section, we obtain an implicit equation for the smallest positive eigen-value *ν*\* for which *C*(*ν*\*) = 0, with *C*(*ν*) given by

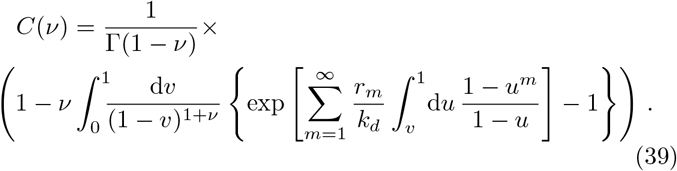

In the limit when ∑*_m_r_m_/k_d_ ≫ 1*, we obtain the following approximate explicit expression for *ν*\*,

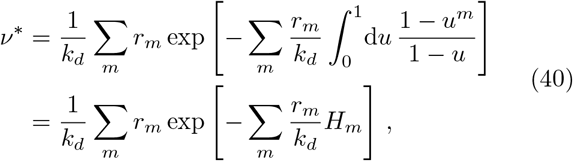

and the corresponding characteristic domain lifetime

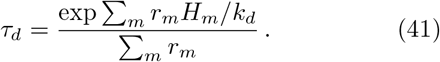

The quasi-stationary distribution of domain sizes is given by the normalized *A_n_*. They can be computed using the recursion relation

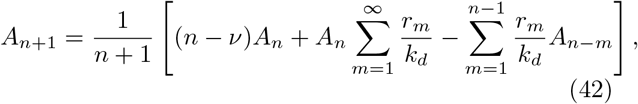

*n* ≥ 1, with *A*_1_ = *ν* ensuring proper normalization. This follows from the fact that domain disappearance only occurs when domains reach size one. Namely, one has

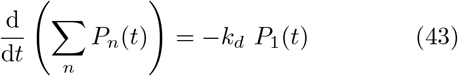

or

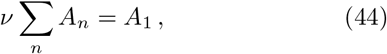

where Eq. (44) holds for a relaxation eigenvector and readily shows that the sum of the *A_n_* equal 1 when *A*_1_ = *ν*. This can also be directly seen by taking *λ* = 1 in Eq. (37).

## III. APPLICATION TO DIFFUSING AND AGGREGATING CLUSTERS

### A. Lifetime and size distribution of an immobile domain: shot-noise appoximation vs. particle-based simulations

In the previous sections, the lifetime of a domain was computed by neglecting correlations between the impinging clusters. In ref. [3], we checked that this type of meanfield approximation produced a cluster size distribution for the diffusing clusters that agreed very well with the results of full numerical simulations. In the present section, we wish to test how these two types of description compare for domain lifetimes.

In the full numerical simulations, single particles and clusters diffuse and aggregate upon encounter. The diffusion coefficient of single particles is *D*_0_ and the clusters diffuse with a size-dependent diffusion coefficient *D_m_*,

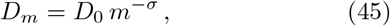

where *m* is the cluster size. The exponent *σ* controls the dependence of diffusion on cluster size. By comparing to experimental data, its value for gephyrin oligomers in spinal cord neurons *ex vivo* was determined to be *σ* ≃ 0.5.

Upon encounter, clusters aggregate and undergo some level of rearrangement. The two extreme cases of no rearrangement and full rearrangement were explored in [3]. No significant effet on the cluster size distribution was observed in the parameter range explored. Here, we limit ourselves, for simplicity, to simulations were clusters are fully rearranged into disks of density *ρ* upon aggregation.

Finally, diffusing single particles and particles in clusters are removed at a rate *k_d_* to mimic turnover. They are reinjected as single particles so that the total concentration of particles in the simulation is constant and equal to *c*_0_. Further simulation details are provided in Appendix C and in ref. [3].

As reported in ref. [3], on an intermediate timescale ~ 1*/k_d_*, the stochastic aggregation-removal dynamics leads to a non-stationary steady state with a characteristic distribution of clusters of sizes *m* with respective concentrations *c_m_*, as shown in Fig. 3A.

**FIG. 3.**
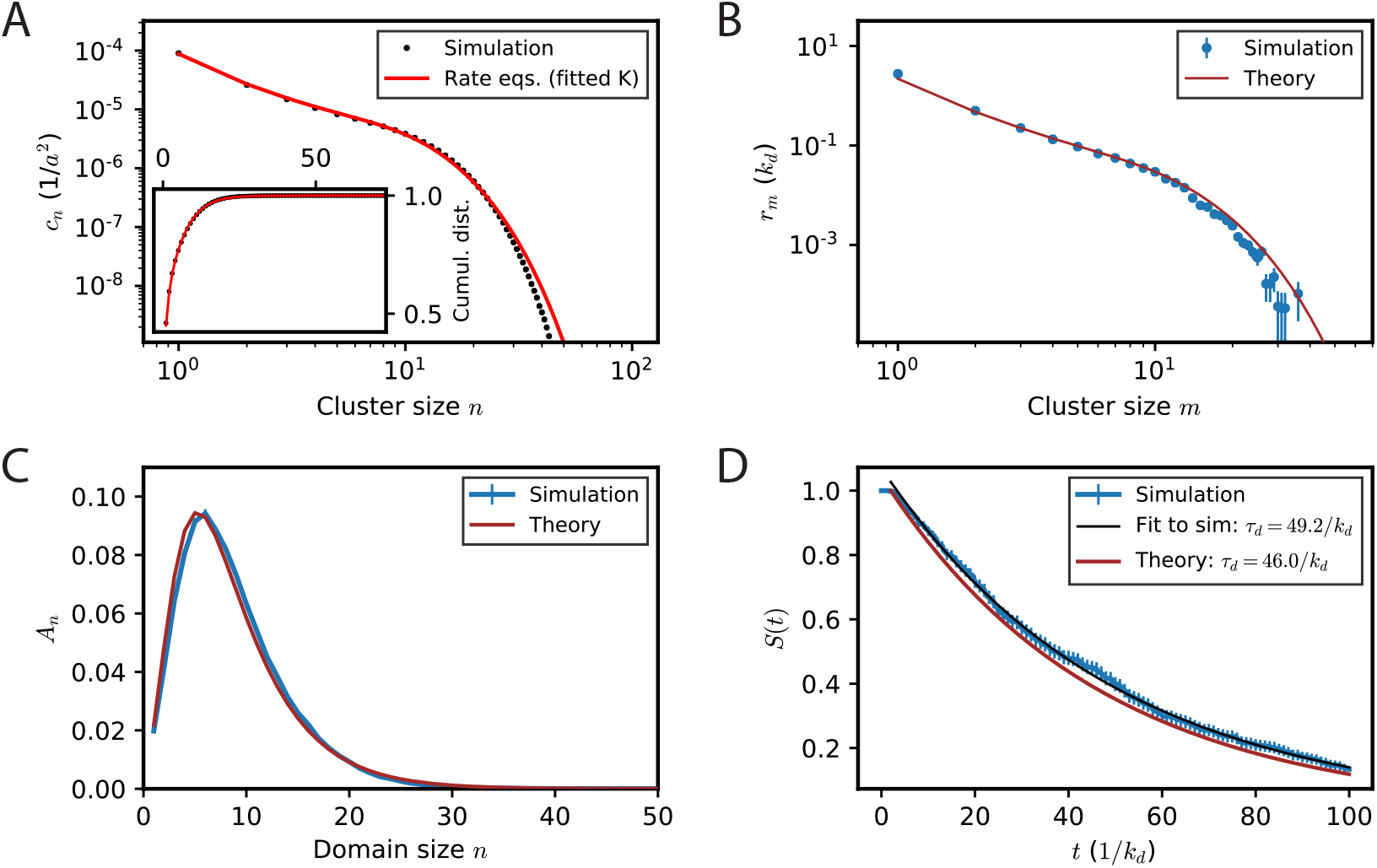
Shot-noise description vs. particle-based simulations of cluster-cluster aggregation for an immobile domain. Simulation parameters are given in Table I, case I. (A) The stationary cluster size distribution (average concentrations of clusters of a given size on the membrane) observed in the simulations (black dots) is shown together with the result from the rate-model description (red line). The latter requires to determine a single free parameter *K* (see text) that is obtained by fitting the cumulative distribution of cluster sizes shown in the inset. (B) The observed rates *r_m_* of encounters with clusters of size *m* for an immobile cluster (blue dots) are shown together with the rate-model approximation (brown line) using the previously determined parameter *K*. (C) Quasi-stationary probability distribution *A_n_* of explored domain sizes for an immobile domain as observed in simulations (blue line) and predicted by the theory with the previously fitted value of *K* (brown line). (D) Clusters disappear due to stochastic fluctuations with a rate 1*/τ_d_*, where *τ_d_* is the typical cluster lifetime. The fraction of surviving clusters *S* observed in simulation (blue) is well-fitted by a decreasing exponential with a time constant *τ_d_* = 49.2*/k_d_* (thin black line). The theoretical prediction based on the rate equations is *τ_d_* = 46.0*/k_d_* (brown line). The approximate analytical expression of the typical lifetime is about 7% shorter than the simulation result.

This dynamics and the steady-state size distribution are well described by the Smoluchowski equations,

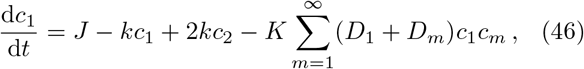

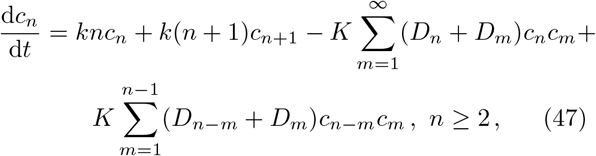

where *J* fixes the total amount of particles *c*_0_ = *J/k_d_*. The kinetic coefficient *K* is determined by fitting the stationary concentrations 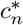 obtained by simulating Eqs. (46,47) to the ones observed in full particle-based simulations, as detailed in Appendix D. The theoretical (Smoluchowski rate equation) result with the fitted parameter *K* is shown in red in Fig. 3A with the cluster distribution obtained from full numerical simulations for comparison.

In order to compute domain lifetimes in full numerical simulations, one particular aggregate of particles, or domain, is initialized with a given number of constituent particles. It is immobilized or fixed during the simulation (i.e. its diffusion constant imposed to be zero) while its size fluctuations and lifetime are monitored over time, see e.g. the blue trajectory in Fig. 1D.

For many parameter choices, the exponential dependence of the domain lifetime on the particle turnover rate *k_d_* and encounter rates *r_m_* renders their lifetime too long to be measured in simulations. Therefore, the simulation parameters (Table I, case I) are chosen in Fig. 3 such as to observe the disappearance of aggregates within reasonable computer time.

These results can be compared with those produced by the shot-noise model of section II. The rates of encounter of a single fixed domain with clusters of size *m* are obtained from the mean-field Smoluchowski description as,

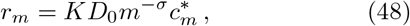

independent of the domain size *n*(*t*). A comparison of these theoretically predicted rates and the ones measured in the particle-based simulations is shown in Fig. 3B. The quasi-stationary distribution of domain sizes is thus readily calculated according to our previous analysis (Eq. (42)), and the characteristic domain lifetime is given by Eq. (41). While the observed cluster size distribution (Fig. 3C) is well described, the predicted characteristic lifetime (Fig. 3D) is ~ 7% off, with *T*_pred_ = 46*/k_d_* vs. *T*_obs_ = 49.2*/k_d_*. This seems rather satisfactory given the exponential dependence on the *r_m_* that involve the fitted parameter *K*. The variation of *T*_pred_ with *K*, to gether with different choices to determine the optimal *K*, is presented in Appendix D.

**TABLE I.** Parameters for computer simulations. In case I, parameters are chosen such that aggregates remain relatively small on average and disappear on timescales compatible with the duration of our simulations, of the order of 100*/k_d_*, see Fig. 3. In case II, we use the parameters identified in a previous work to describe the aggregation-removal dynamics of gephyrin domains in spinal cord neurons [3]. The quasistationary domain size distribution in that case is shown in Fig. 5. Note that parameters are non-dimensionalized using the particle density in aggregates *ρ* and the diffusion constant for monomers *D*0. See ref. [3] for details on the implementation.

|  | case I<br>(fast turnover) | case II<br>(slow turnover) |
| --- | --- | --- |
| $c_0/\rho$ | $10^{-3}$ | $0.9 \cdot 10^{-3}$ |
| $k_d/(c_0 D_0)$ | 0.1 | 0.022 |
| $\sigma$ | 0.5 | 0.5 |

In Fig. 3D, the observed domain lifetime is determined by fitting an exponential decay to the fraction of surviving domains. As a check for internal consistency, we can furthermore compare this result with the observed probability of domains to be monomeric, i.e., the observed *A*_1_. Within the precision of our simulations, the measured lifetime *T*_obs_ is consistent with the expected lifetime 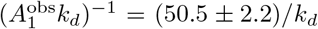, where the uncertainty corresponds to one standard deviation determined by non-parametric bootstrap resampling from different realizations of the simulation (see Appendix C for details).

### B. Lifetime and size-distribution of a diffusing cluster

In the previous subsection as well as in section II, we focused on the size fluctuations of an immobile domain surrounded by diffusing and aggregating clusters, which appears the most interesting case in the context of synaptic dynamics. Nonetheless, one can also imagine following a diffusing cluster of particles over time. As the cluster diffuses, it encounters other diffusing clusters which makes its size grow, while losing single particles by turnover, see e.g. the orange trajectory in Fig. 1D. The size distribution obtained by following a diffusing cluster over time is similar to that of a fixed domain, see Fig. 1E, but very different from the size distribution of diffusing clusters at a given time instant (Fig. 1C). Namely, when a cluster is followed, it is observed that irrespective of its initial size, after a time of a few 1*/k_d_*, it fluctuates over a mean average size in a quasi-stationary fashion [3].

One can try and adapt the methods developed for immobile domains to determine the distribution of sizes that a diffusing cluster explores over time, as well as its average lifetime.

A simplified shot noise process for a diffusing cluster can be written as for an immobile domain. The corresponding master equations generalizing Eq. (36) read

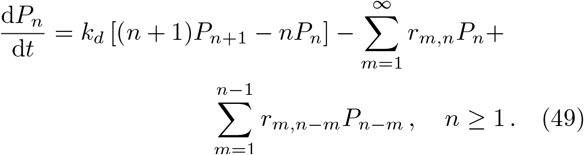

When the diffusion constant of the clusters is given by Eq. (45), the rates *r_m,n_* are given by

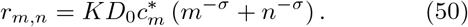

The difference with the case of an immobile domain (Eq. (48)) is that the rates *r_m,n_* do not only depend, as previously, on the size *m* of the impinging clusters but also, when *σ* ≠ 0, on the size *n* of the followed diffusing cluster. A comparison between the rates given by Eq. (50) and measured encounter rates (see Appendix C for details) in full particle-based simulations is shown in Fig. 4A.

**FIG. 4.**
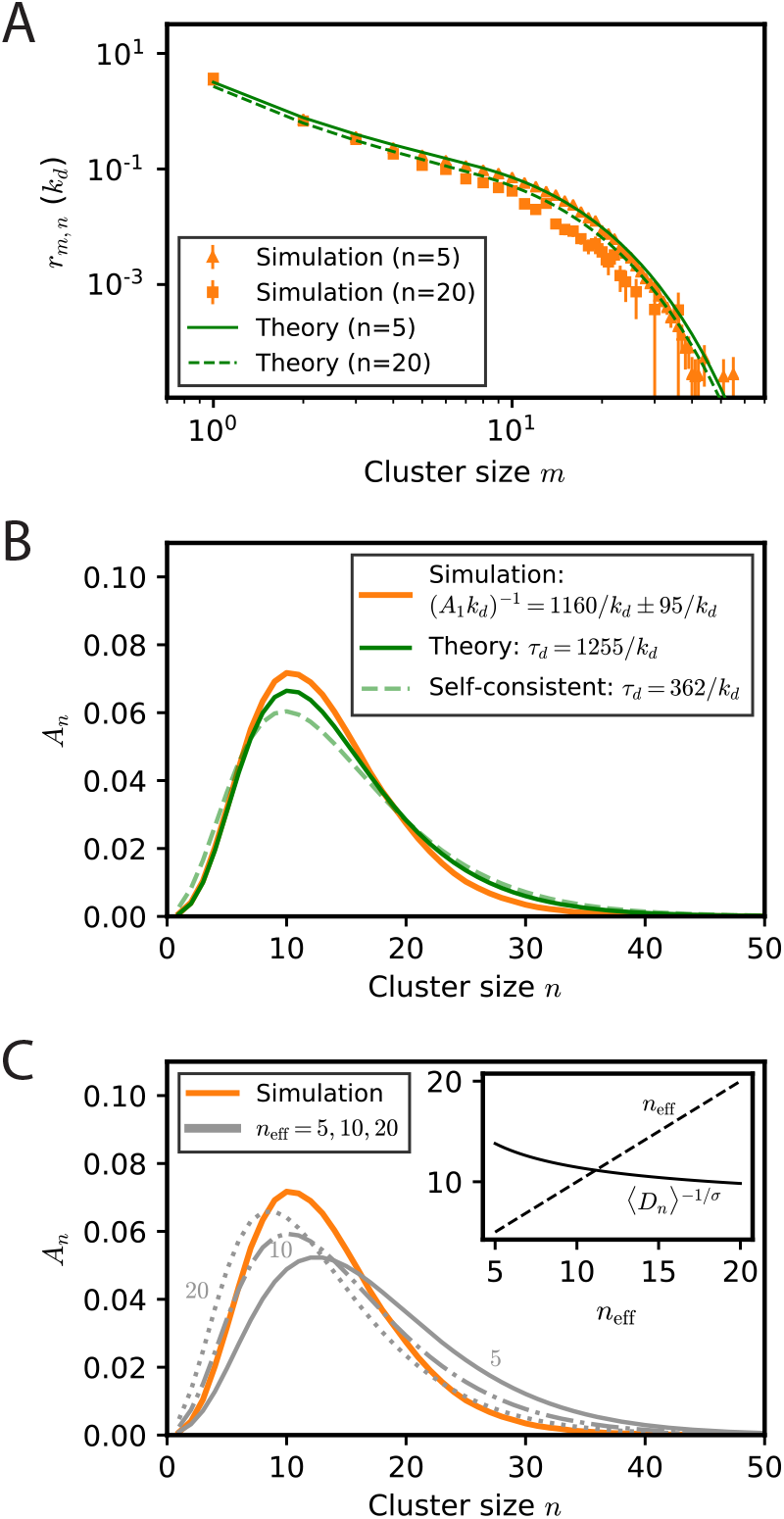
Shot-noise description vs. particle-based simulations of cluster-cluster aggregation for diffusing clusters. Simulation parameters are given in Table I, case I. (A) The observed rates *r_m,n_* of encounters of a cluster of size *n* (orange triangles: *n* = 5; orange squares: *n* = 20) with clusters of size *m* are shown together with the rate-model approximation using the previously determined parameter *K* (Fig. 3A). (B) The quasi-stationary probability distribution of explored cluster sizes for freely diffusing clusters as observed in the simulations (orange line) and predicted (solid green line) from the rate-equations with full size-dependence of encounter rates *r*_*m,n*_ (Eq. (50)) and choosing the value *ν* = *ν*\* which gives a fast decay of the *A_n_* for large *n*. The theoretically predicted probability distribution obtained with a self-consistently determined effective cluster size *n*_eff_ and corresponding encounter rates *r_m_* that do not depend on the size *n* of the tracked cluster is also shown (dashed green line). The respective predicted characteristic lifetimes are given in the plot legend. The given standard deviation is obtained using the bootstrap method explained in Appendix C 3. (C) Determination of the self-consistent cluster size *n*_eff_ . The predicted quasi-stationary probability distribution *A_n_* of observing a diffusing cluster of size *n* depends on the effective cluster size *n*_eff_ via the effective encounter rates; distributions are shown for *n*_eff_ = 5 (solid gray line), *n*_eff_ = 10 (dashed-dotted gray line) and *n*_eff_ = 20 (dotted gray line); the distribution obtained from the simulation is also plotted (orange line, same as in panel B). For larger *n*_eff_, the rates are lower and the size distribution tends to lower *n*. Inset: graphical explanation of the self-consistent determination of *n*_eff_ . The expected average cluster diffusion constant ⟨*D_n_*⟩ = ∑_*n*_ *A_n_*(*n*_eff_)*D_n_* depends on *n*_eff_ via the *A_n_* and increases with *n*_eff_ . A self-consistent solution for the effective size is then given by *n*_eff_ = (⟨*D_n_⟩/D*_0_)^−1*/σ*^, obtained as the crossing of the curve representing the expression of the r.h.s. (solid line) with the diagonal (dashed line).

As discussed above, the quasi-stationary distribution and the lifetime of the diffusing cluster correspond to the eigenvector of Eq. (49), *P_n_* = *A_n_* exp(−*νk_d_t*), with the smallest decay time, i.e. the smallest possible *ν*. Substitution into Eq. (49) allows one to recursively determine the *A_n_*,

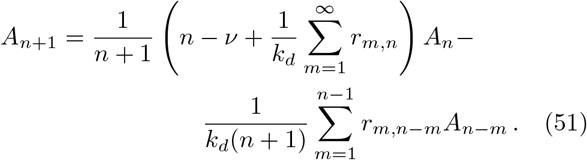

The desired eigenvalue is obtained as the value of *ν* = *ν*\* for which the prefactor of the slow decay Eq. (26) vanishes. The corresponding quasi-stationary distribution of the sizes *n* explored by a diffusing cluster is given by the corresponding normalized *A_n_*.

The lifetime and quasi-stationary distribution computed from this analysis of the model Eq. (49) are compared to full numerical simulations of diffusing and aggregating clusters in Fig. 4B for the same parameters as those of Fig. 3. The shot-noise description is found to account for the particle-based simulations similarly well for diffusing clusters as for an immobile domain.

For an immobile fluctuating domain, we could compute numerically the eigenvalue *ν*\* but also obtain the analytic expression (40) for the function *C*(*ν*) of which *ν*\* is a root, as well as the explicit asymptotic expression (39) for *ν*\* in the parameter regime in which domains are long-lived. When the rates *r_m,n_* explicitly depend on the size of the domain considered, these analytic computations appear difficult without further approximations. We did not succeed in finding an analytic solution for the integro-differential equation that replaces the differential equation (37) for the generating function of the *A_n_*.

One simple approximation that we tried is to replace the rates *r_m,n_* by rates 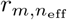 where the varying size of the considered cluster has been replaced by a fixed effective size *n*_eff_. This eliminates the additional size dependence of the rates on the size *n* of the diffusing cluster and reduces the calculation to that performed for an immobile domain. The resulting quasi-stationary distributions *A_n_*(*n*_eff_) for different values of *n*_eff_ are shown in Fig. 4C for the same parameters as those of Fig. 4B. One can moreover try and determine *n*_eff_ self-consistently by requiring for instance that the mean cluster diffusion constant obtained from the distribution *A_n_*(*n*_eff_) be equal to the effective diffusion constant that served to determine the distribution,

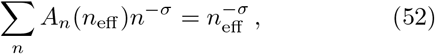

where we have used the expression (45) for the diffusion constant. The self-consistent determination of *n*_eff_ obtained from Eq. (52) is depicted in the inset of Fig. 4C. As shown in Fig. 4B, the resulting quasi-stationary distribution provides a fair approximation of the exact quasistationary distribution for Eq. (49). However, the diffusion rate of the cluster is underestimated in the self-consistent approximation when the cluster size is smaller than *n*_eff_ . This gives longer estimated residence times of the cluster at small sizes and a larger weight at *n > n*_eff_ for the self-consistent quasi-stationary distribution than for the exact one. This discrepancy leads the the selfconsistent approximation to significantly underestimate the diffusing cluster lifetime (by a factor of about 3.5 for the parameters of Fig. 4). More accurate approximations of the quasi-stationary distribution for small cluster sizes would thus be worth developing.

### C. Lifetime of postsynaptic gephyrin clusters

In ref. [3], we determined biophysical parameters governing the aggregation-and-removal dynamics of postsynaptic gephyrin aggregates. For representative values (see Table I, case II), the size distribution of diffusing clusters obtained in particle-based simulations is shown in Fig. 5A. It is well reproduced by the distribution produced by the mean-field Smoluchowski equations (46,47)) with the parameter *K* fitted using the cumulative cluster size distribution (Fig. 5A insert), as explained in Appendix D. As shown in Fig. 5B, the impinging cluster rates on the immobile fluctuating domains obtained from the Smoluchowski equations also agree well with the rates directly measured in particle-based simulations.

**FIG. 5.**
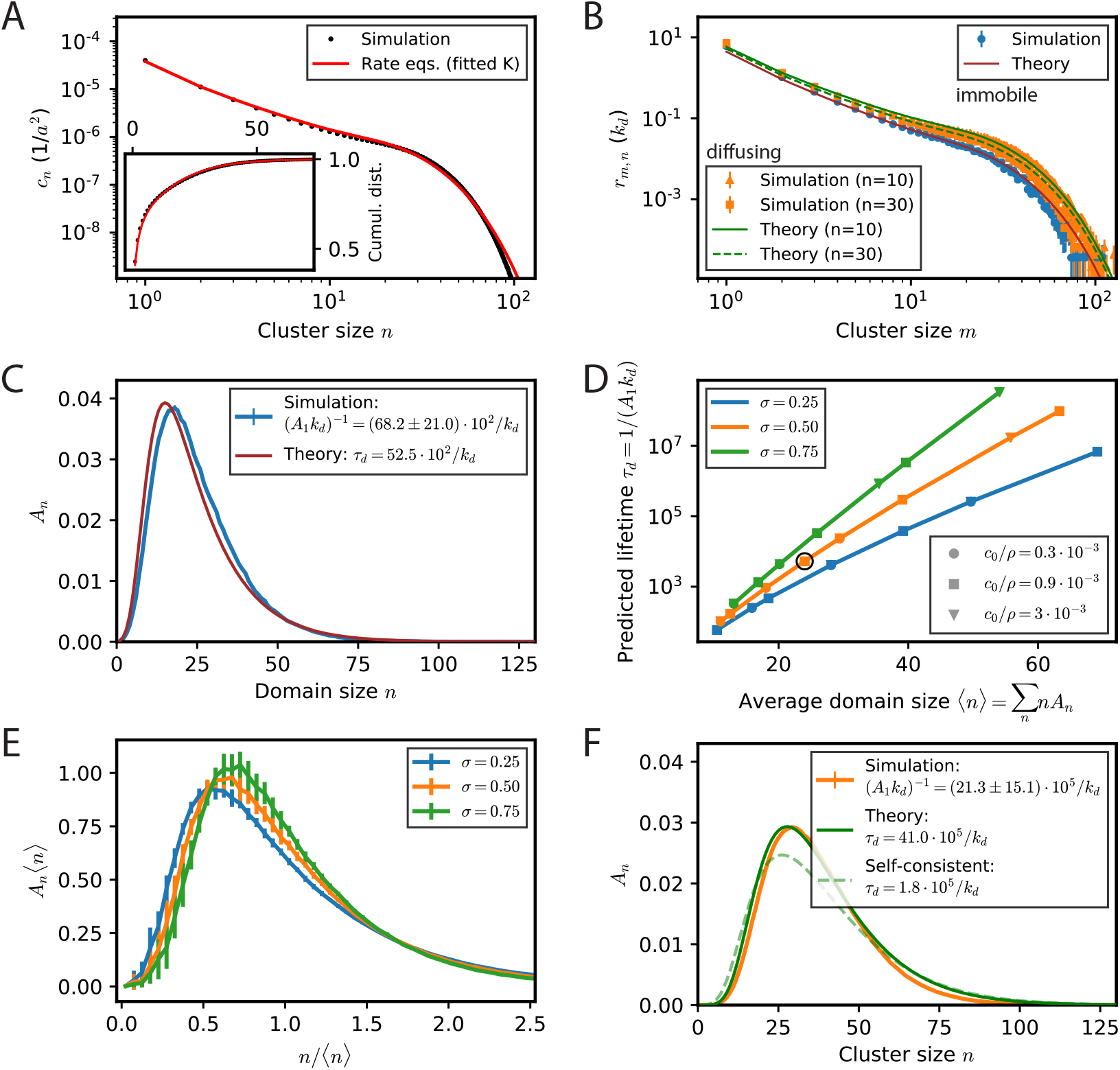
Predicted distribution of explored cluster sizes for synaptic gephyrin domains, which allow to infer the typical domain lifetime. Simulation parameters are given in Table I, case II. (A) Particle-based simulations with the parameter values obtained in a previous work [3] give rise to a stationary distribution of diffusing clusters in the membrane (black dots). The stationary cluster size distribution (average concentrations of clusters of a given size on the membrane) is shown together with the result from the rate-model description (red line). The latter requires to determine a single free parameter *K* (see text). Inset: The best fit to the cumulative distribution of cluster sizes is obtained for *K* = 1.81. (B) The observed rates of encounter *r_m_* for domains of size *n* with clusters of size *m*. For immobile domains (blue dots), the rates do not depend on the size *n*; for diffusing clusters that are followed, the rates are shown for *n* = 10 (orange triangles) and *n* = 30 (orange crosses). The theoretical predictions for the *r_m_* based on the fitted *K* (panel A) are shown as lines (immobile: solid brown, diffusing *n* = 10: solid green, diffusing *n* = 30: dashed green). (C) The quasi-stationary distribution of sizes explored by immobile domains subject to encounter with diffusing clusters and turnover of constituent particles for particle-based simulations (blue line) and the prediction from the rate-equation description with fitted *K* (brown line). The expected lifetime of immobile domains is given in the plot legend, where the uncertainty in the simulation result reflects limited sampling of occurrences where the domain size is 1. (D) Theoretically predicted lifetimes for immobile domains for different values of *c*_0_, *k_d_*, and *σ*. For given *σ*, the predicted lifetime depends only on the average domain size *n* . The parameters corresponding to the simulations and theory shown in panels A-C,F are surrounded by the black circle; for all parameters, we used the value *K* = 1.81 for the mean-field equations as determined in panel A. (E) The averaged rescaled quasi-stationary size distributions of immobile clusters for different combinations of *c*_0_, *k_d_*, and *σ* show a trend towards narrower size distributions with increasing *σ*, implying larger domain lifetimes. Error bars represent the standard deviation over the averaged quasi-stationary size distributions for different *k_d_* and *c*_0_. (F) For diffusing domains, the distribution of explored cluster sizes tends to larger values as the rates of encounter with other clusters are larger on average than for immobile domains, see panel B. The self-consistent solution (see text for details) captures the overall shape of the distribution but can be expected to overestimate the probability of observing clusters of size 1, see text for details. Because the corresponding cluster lifetimes are extremely long, the quantitative accuracy of the predictions is difficult to assess by direct numerical simulations.

Using these rates, we can compare the predicted quasistationary size distribution of surviving clusters to the one observed in particle-based simulations, see Fig. 5C. In our simulations, fixed clusters are on average composed of 25 ± 13 gephyrin trimers (± standard deviation), which matches well the analytical prediction 24 ± 14. (Here, we consider a single particle to represent a gephyrin trimer, as the trimeric form of gephyrin is very stable and gephyrin does not seem to exist as a monomer or in dimeric form in cells.) Because of the relatively large cluster sizes involved and slow particle loss, the expected typical cluster lifetime is much longer than for the other parameters studied, and therefore difficult to observe in simulations. Our analytical result Eq. (41) nevertheless provides us with a quantitative estimate: For gephyrin domains whose dynamics are governed by the processes described above we predict an average lifetime of *T*_pred_ ≈ 5.25 · 10^3^*/k_d_*. With a measured *k_d_* of about half an hour [5], this gives a lifetime of 3-4 months. This is several orders of magnitude larger than the turnover time of individual proteins and highlights the ability of the aggregation-removal dynamics to maintain (on average) stable synaptic domains over long times. Note that the precise predicted value depends on the value of *K* used for approximating the full dynamics by Smoluchowski rate equations. However, comparison with the expected rate of decay as predicted by the sampled frequency of clusters being of size 1, Fig. 5C, shows both values are consistent within the error margin of our simulations, see also Appendix D.

In ref. [3], fitting the model to experimental distributions actually resulted in a range of admissible values for the concentration of diffusing *c*_0_ of diffusing gephyrin trimers, turnover time *k_d_* and diffusion exponent *σ* (Eq. (48)). A variation in these values results in a change of the rates *r_m_* in the Smoluchowski description. The corresponding variation of the domain lifetime can be simply computed using the shot-noise model. As shown in Fig. 5D, the resulting variation in domain lifetime is large and can be ascribed to the variation in the average size of the fluctuating domain. As expected, the larger the average size of the domain, the longer its lifetime. Domains composed of 70 gephyrin trimers could theoretically have an extremely long lifetime.

There is also a significant influence of the exponent *σ*, with a larger *σ* producing a longer lifetime for the same average domain size. The influence of the exponent *σ* on the quasi-stationary distribution is milder, as shown in Fig. 5E.

A similar analysis can be repeated for diffusing clusters. The theoretically predicted quasi-stationary size distribution is compared with the one determined in simulations in Fig. 5D for parameters of case II (see Table I). The expected lifetime of diffusing domains is several orders of magnitudes larger than for immobile domains. Because the observation of clusters of size 1 is a very rare event, the quantitative accuracy of the theoretical prediction is however difficult to assess. Similarly to the parameters with faster turnover (Fig. 4), the self-consistent solution overestimates small and large cluster sizes because encounter rates are underestimated when the cluster is small and overestimated when the cluster is large.

## IV. DISCUSSION AND CONCLUSION

In this work, we have derived an explicit expression for the lifetime of a mesoscopic domain of particles that evolve, i.e., respectively grow and shrink, by the aggregation of clusters of particle of different sizes and loss of individual constituent particles. This was achieved by considering the simpler mathematical problem of computing the average time of first passage to zero for a Markov process with downward jumps of unit size, representing particle loss, and upward jumps of a range of sizes, representing cluster aggregation, at rates only depending on the size of the jumps, i.e., simply on the size of the impinging clusters and their concentrations. This is a simplification of the spatial model of ref. [3] in the spirit of the mean-field Smoluchowski equations [22]. These are known to give an accurate description of the cluster size probability distribution [23] after the adjustment of an overall kinetic parameter, as it was also checked in [3]. Similarly, we have found here that our derived expression for the domain lifetime holds approximately for full particle-based simulations of the physical processes of diffusion, aggregation, and turnover. This has thus allowed us to provide an estimate of the expected lifetime of inhibitory postsynaptic scaffold domains based on our earlier proposed model [3] together with the previously determined biophysical parameters for diffusion, aggregation kinetics, and turnover of gephyrin scaffolding proteins. Outside of the synaptic domain context, the method and techniques should be generalizable to other cases of interest [14–16] where mesoscopic structures arise through a balance of diffusion-mediated aggregation and recycling.

One motivation of our work was the fundamental question of long-term memory maintenance while the turnover time of synaptic components is on the scale of hours and their typical lifetime on the scale of days [24] (although some synaptic proteins have been found to be very long-lived [25, 26]). A simple idea already proposed by Crick [2] is that the biophysical substrate of longterm memory lies in a multi-particle state, the existence of which could be much longer than the residence time of single particles. Crick himself suggested a cooperative switch based on phosphorylation [2], an idea much further elaborated by Lisman and others who suggested the autophosphorylation of CaMKII as its implementation. While there is strong evidence that CaMKII is involved in early long-term potentiation of synapses, its involvement in their long-term maintenance is much less clear since CaMKII catalytic activity only persists for about minute after synaptic stimulation (see [27] for a review).

A long-lived multi-particle state is also at the root of a different proposal by Shouval [28] who suggested that clustering of neurotransmitter receptors at the synapse increased the insertion rate of new receptors. This allows for a clustered state of many receptors to persist at the synapse while individual receptors go in and out on a much shorter timescale. In the present work, our emphasis has been on the role and dynamics of synaptic scaffolding proteins rather than on those of receptors. We have shown that a domain persistence time much longer that the residence time of individual scaffolds naturally occurs in the model of ref. [3] for an oligomer of scaffold proteins similar to those found at inhibitory synapses.

The model of ref. [3] can be experimentally distinguished from other ones [28–30] on different grounds. First, at the most basic level, it supposes that the existence of a scaffold domain depends on a lateral flux of scaffolding proteins onto the domain. This can be experimentally tested by perturbing this flux and assessing the resulting effect on the domain. Second, the model predicts specific size distributions of diffusing clusters. Extrasynaptic scaffold clusters are seen experimentally [3, 8]. The measured distributions obtained in a previous work are consistent with the model predictions [3] but more data could be gathered for more precise tests. Third, the temporal fluctuations of domain size analyzed in the present work are a specific prediction of the model and their measure should be experimentally accessible.

Our description is admittedly a provocatively simplified description of a synapse, in different respects. First, we have not considered the presynaptic side, or only assigned to it an implicit role in helping to fix the position of the postsynaptic domain considered. Second, the postsynaptic side is known to harbor many other proteins besides receptors and scaffolds, such as adhesion, cytoskeletal and signaling proteins. These proteins as well as the interaction between the preand postsynaptic sides certainly play a role on synapse lifetime and modify our estimate. Nonetheless, the present study highlights that the proposed dynamics of postsynaptic scaffold domains already endows them with a much longer lifetime than those of individual scaffold domains. A long lifetime of synaptic domains is thus compatible with a turnover of scaffold proteins on the hour time scale. One specific assumption we made is that the lifetime of a domain is determined by its shrinkage to a vanishing size. It is certainly worth trying to examine this scenario experimentally and more generally to experimentally investigate the path of a synapse towards disappearance.

Besides lifetime *per se*, the explicit model we have examined highlights features of synaptic dynamics that need to be reconciled with the function of synapses in memory maintenance. Namely, our model synaptic domain is subject to large fluctuations (see Fig. 1D&E and 5) so that correlations of its size between different instants in time quickly decay with their distance in time (Eq. (7)). The probability that the synaptic domain has a given size at time *t* is simply described by the quasistationary size distribution, independently of the size it had a few turnover times before, as it has also been seen experimentally, both for excitatory [31, 32] and inhibitory synapses [29] (although with a slower relaxation rate than in the present model). This is also seen in other proposed models of synaptic fluctuations e.g. [30]. This emphasizes the much greater difficulty of preserving in a persistent way the size of a domain or the “strength” of synapses than maintaining their mere existence. This is a general difficulty for the encoding of a continuous variable, which inevitably tends to drift in the presence of fluctuations. Future work will hopefully tell us whether longterm memory maintenance relies on the use of multiple synapses between neurons, or synapses with discrete values, perhaps based on synaptic nanodomains [33], or in another yet unforeseen manner.

## ACKNOWLEDGMENTS

We wish to thank Antoine Triller and Christian Specht for many instructive discussions on inhibitory synapses.

## Appendix A: Large-*n* asymptotics of the stationary and quasi-stationary distributions

The asymptotics of the stationary probability distribution *P_n_* (Eq. (13,14,15)) and the quasi-stationary one *A_n_* (Eq. (23,24)) can be obtained either from their generating functions or directly from the recurrence relations that they obey, as we briefly discuss.

We take the stationary probability distribution *P_n_* for monodisperse diffusing clusters of size *m* as an example. The generating function *P* (*λ*) for the *P_n_* is defined in Eq. (16). Its large-*λ* behavior is directly obtained from the explicit expression (Eq. (18)) of *P* (*λ*) as,

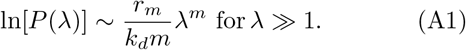

From the definition of *P* (*λ*) (Eq. (16)), the method of stationary phase gives that the asymptotics (A1) for *λ* ≫ 1 corresponds to the size *n* that maximizes *g*(*y*) = *ny* + ln(*P_n_*), namely to the Legendre transform of − ln(*P_n_*) with respect to the variable *y* = ln(*λ*). The logarithm of the probability ln(*P_n_*) can thus be obtained by taking the extremum of −*ny* + *g*(*y*) with respect to *y*,

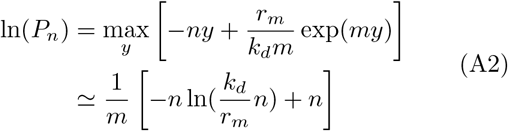

Further terms in the asymptotics of *P_n_* could be obtained by matching the expansion around the stationary phase to the large *λ* asymptotics of the generating function *P* (*λ*). It is also possible to obtain them directly from the defining recursion relations *P_n_* as we now show.

For *n > m*, the *P_n_* obey (Eq. (14)),

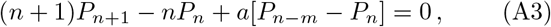

where we have defined *a* = *r_m_/k_d_* to simplify notations. For large *n*, some terms dominate and their balance determines the asymptotics. The balance of the first two terms in Eq. (A3) is a simple possibility. It gives *nP_n_ ~ cst* which is consistent in the sense that the two neglected terms proportional to *a* are indeed found to be negligible with respect to the kept terms. However, this slow asymptotics gives a non-normalizable function which cannot correspond to the distribution *P_n_*. Another possibility is that *P_n_* decreases sufficiently quickly with *n*, so that *P_n_*_−*m*_ is comparable to *nP_n_*. The balance of the two corresponding terms in Eq. (A3) gives back very simply the previously found asymptotics (Eq. (A2)),

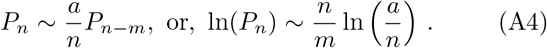

The fast asymptotics (A2) or (A4) can be obtained more systematically by searching *P_n_* in an exponential WKB-like form [34],

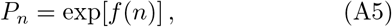

with a slowly varying *f* so that its value at *n − m* can be approximated by a Taylor expansion around *n*. With the help of Eq. (A5), the recursion relation (A3) can be rewritten as

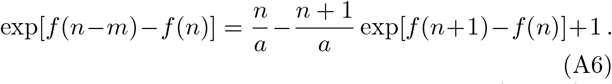

After expansion of *f* in Taylor series, e.g. *f(n − m) = f(n) − mf′(n) − m^2^f″(n)/2+ …* one obtains at dominant order

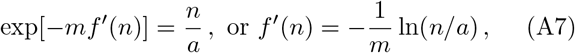

which is equivalent to the previously found dominant asymptotics of *P_n_* (Eq. (A2)). The first few subdominant asymptotic terms in *f* (*n*) can be obtained by searching for *f* ′(*n*) under the form,

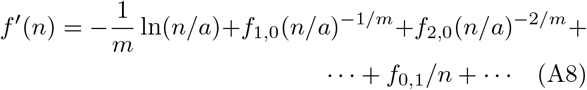

where the dots denotes terms with mixed negative powers of *n*^1*/m*^ and *n* and eventually other more complicated subdominant terms. Substitution in Eq. (A6) after Taylor expansion of the finite difference terms determines the coefficients *f*_10_*, f*_2,0_ and *f*_0,1_,

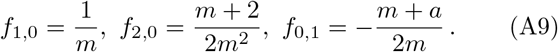

Finally, this provides by integration the asymptotic series for *f* (*n*) which refines Eq. (A2),

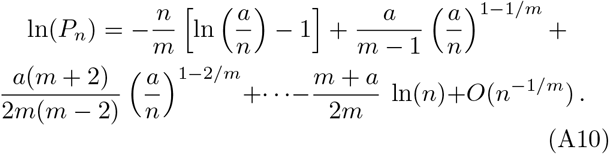

## Appendix B: Local expansions and explicit expression for *ν*\*

We detail here our derivation of the local expansion (30) and of the expression (35) for the root *ν*\* of the function *C*(*ν*).

In order to obtain Eq. (30), we write the expression (29) for the generating function *A*(*λ*) as

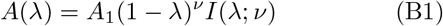

with

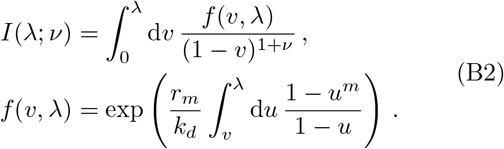

In the limit of *λ* → 1, we can develop the integrand and obtain to first order

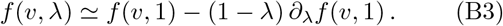

Because the second term scales with an additional power (1 − *λ*), it does not contribute to the two leading terms in the expansion of *I*(*λ*; *ν*) around *λ* = 1, *I*(*λ*; *ν*) = *ν*^−1^(1 − *λ*)^−ν^ + *B*(*ν*) + …, and we can safely neglect it.

As *f* (*υ*, 1) is finite over the range of integration, the integral *I*(*λ*; *ν*) is dominated by the divergence at its upper bound *υ* = *λ* for *λ* → 1. We therefore split the integral in two parts, *I* = *I*_1_ + *I*_2_, with

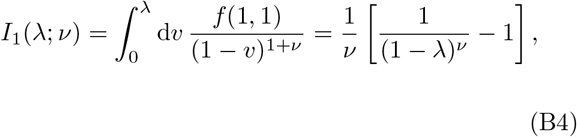

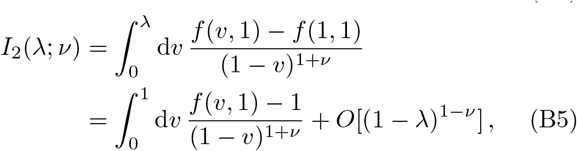

where we have used *f* (1, 1) = 1 in the second equalities of Eq. (B4) and (B5). Note that *I*_2_(1; *ν*) does not diverge because *f* (*υ*, 1) − *f* (1, 1) vanishes like *v* − 1 as *v* → 1. Substitution of *I* by the expressions (B4) and (B5) for *I*_1_ and *I*_2_ provides the local expansion (30) and the explicit expression (34) for *C*(*ν*).

In the limit when *a* ≡ *r_m_/k_d_* is large, *I*_2_(1; *ν*) becomes independent of *ν* and the root *ν*\* of *C*(*ν*) is readily obtained. From the expression of *f* (*υ*, 1) (Eq. (B2)), it is clear that its maximum of the integrand for *I*_2_(1; *ν*) is at *υ* = 0. For *a* ≡ *r_m_/k_d_* ≫ 1, the dominant contribution to the integral comes from a neighborhood of *υ* = 0 of size 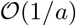 since

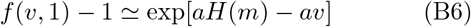

where the number *H*(*m*) is given by Eq. (21). In this small neighborhood, the denominator of the integrand is simply equal to one to dominant order, (1 − *λ*)^1+*ν*^ ~ 1, and

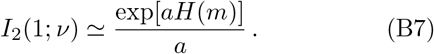

Replacement of *I*_2_(1;*ν*) by this *ν*-independent approximation in the expression for *C*(*ν*) (Eq.(34)) immediately provides the explicit expression (35) for *ν*\*.

## Appendix C: Numerical simulations

### 1. Simulation of the shot-noise process

To simulate the shot-noise process described and analyzed in section II, we implemented a discretized version of the dynamics described by Eqs. (13,14). For each time step *t_i_*, the domain size *n*(*t_i_*) was diminished by 1 with probability *n*(*t_i_*)*k_d_*∆*t* and increased by *m* with probability *r_m_*∆*t* to determine *n*(*t_i_*_+1_). The domain lifetime was determined as the first passage to *n* = 0, where domain sizes were initialized at the expected mean cluster size ⟨*n*⟩ = *r_m_m/k_d_*. In our simulation, ∆*t* = 0.001 < 1*/*(⟨*n⟩k_d_*), with *k_d_* = 1, proved sufficiently small to show excellent agreement between simulations and analytic results. We simulated the size trajectories of 2000 domains over a duration of *T*_sim_ = 500*/k_d_*.

### 2. Numerical determination of *C*(*ν*)

We used the large-*n* asymptotics Eq. (26) to determine 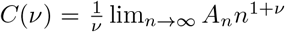 from the corresponding recursion relation in the case of mono- or polydisperse aggregation for immobile domains (Eqs. (23,24) or (42), respectively) or for diffusing domains (Eq. (51)). For practical reasons, we chose a large, finite *n_c_* to evaluate 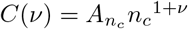 and checked that *C*(*ν*) did not significantly vary with increasing *n_c_*.

### 3. Full spatial particle-based simulations

The full numerical simulations reported in the present article correspond to those described as model A in ref. [3]: upon aggregation or single particle loss, a cluster is immediately rearranged into a disc of radius 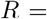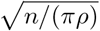, where *n* is the number of particles in the cluster and *ρ* the cluster density. Simulations are performed as described in ref. [3].

The diameter of a single particle *a* and the diffusion time *a*^2^*/D*_0_, are used as unit of length and time, with *D*_0_ the diffusion constant of a single particle. Time is discretized in steps of ∆*t* = 0.02. Space is a square box of side length *L* with periodic boundary conditions. The cluster density was chosen to be *ρ* = 0.77*a*^−2^ and corresponds to the packing fraction of a hexagonal lattice *ρ* = 0.77*a*^−2^.

A simulation consists of the simulated dynamics of *N* = *c*_0_*L*^2^ particles. In simulations with an immobilized domain, a cluster of size *n*_init_ is initially positioned at the center of the simulation box and its diffusion constant set to zero throughout the simulation, including after fusion events with impinging clusters. The remaining *N − n*_init_ particles are randomly positioned. In simulations without an immobilized domain, all *N* particles are initially positioned at random. Then, at each time step, the four following moves are performed. First, particle and cluster positions are displaced by random Gaussian increments corresponding to their diffusion constants. Second, overlapping particles or clusters are merged and their radius updated. Third, the number of removed particles per cluster is drawn from a binomial distribution and the cluster sizes and radii are reduced accordingly. Fourth, removed particles are reinserted as new randomly distributed particles. The total number of particles in the simulation is thus constant.

In the present article, two different parameter sets were used for the simulations, see Table I. In addition, we used *L* = 1000 and *n*_init_ = 10 (where appropriate) for parameter set I (Figs. 3 & 4) and *L* = 2000 and *n*_init_ = 20 (where appropriate) for parameter set II (Figs. 1 & 5). All simulations were run for a total time of *T*_sim_ = 100*/k_d_*.

Reported quantities were measured as follows: the instantaneous cluster size distribution *c_n_* was determined as the average over realizations and time after a transient of 10*/k_d_*. The explored cluster size distribution was determined from individual size trajectories of immobile or diffusing clusters, taking into account all sizes after an initial transient of 10*/k_d_* until the (potential) disappearance of the cluster. For immobile clusters, the lifetime was simply given by the time at which the cluster disappeared; the fraction of surviving clusters is the number of simulations for which the immobile domain still exists relative to the total number of realizations. To determine the encounter rates, all individual fusion events in the simulations were registered. (In the case of immobile domains, only fusions with the immobile domain where registered.) For the immobile domains, the encounter rate was given by the number of fusions with clusters of a given size *m* divided by the time over which fusions were tracked. For diffusing domains, all events where clusters of sizes *m* and *n* fused were independently counted and divided by the time over which fusions were tracked; in order to obtain encounter rates that do not scale with the size of the simulation box, these rates were normalized by the expected average number of clusters per size *n*, *N_n_* = *c_n_L*^2^, using the average cluster concentrations *c_n_* observed in the simulations in order to obtain the rates *r_m,n_*. (Fusions were also considered only after a transient of 10*/k_d_*.)

To estimate the uncertainty of simulation results due to finite sampling, we used a non-parametric bootstrap method. When *n*_real_ data points were obtained by simulations, we produced *n*_bootstrap_ synthetic realizations (*n*_bootstrap_ = 1000) by randomly resampling *n*_real_ times these data points with replacement. Where simulation results are reported, errors (respectively error bars in the figures) represent the standard deviation of the distribution of measured values obtained from these *n*_bootstrap_ resampled data. For immobile domains, we analyzed *n*_real_ = 500, respectively *n*_real_ = 300, independent size trajectories for parameter cases I and II, respectively, from the same number of independent simulations. For diffusing domains, we analyzed *n*_real_ = 1557 independent (non-overlapping) cluster size trajectories from 50 independent simulations that lasted at least 20*/k_d_* in case I; in case II, we analyzed *n*_real_ = 4365 independent (nonoverlapping) cluster size trajectories from 100 independent simulations that lasted at least 20*/k_d_*.

## Appendix D: Comparison with Smoluchowski equations: fit of the kinetic coefficient *K*

In order to compare the Smoluchowski description with full particle-based numerical simulations, one has to determine the kinetic coefficient *K* in Eqs. (46,47). This determines the rates *r_m_* (Eq. (48)) and allows one to compute the fluctuating domain lifetime 1*/*(*ν*\**k_d_*) predicted from the implicit equation *C*(*ν*\*) = 0 for *ν*\* with Eq. (39), or from Eq. (41) being the approximate expression in closed form.

We can use the theoretical predictions for the steadystate cluster concentrations 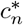 to determine the value of *K* that best accounts for the data, as it is the sole free parameter in the mean-field description for the diffusion aggregation dynamics for known *c*_0_, *k_d_* and *σ*. In a prev ious work [3] we used the typical cluster size 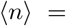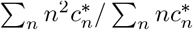 to determine an approximate expression for *K* = *K*(*c_0_*) that best accounted for the ensemble of parameter sets *c*_0_, *k_d_*, and *σ* tested. The precise value of *K* does not matter much as long as only size distributions are concerned, for small variations of *K* translate into only small variations of the predicted cluster concentrations 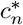. However, because of the exponential dependence of the *τ_d_* on the encounter rates *r_m_* (Eq. (41)), the precise value does matter when exact domain lifetimes are to be predicted.

Here, we tried to determine the *K* that best accounts for the particle-based simulations independently for the two parameter sets investigated. We furthermore compared different ways to determine the optimal *K* in order to address the uncertainty linked to the Smoluchowski mean-field approximation. In all three cases, we determined the optimal *K* by minimizing the least-squares error ∑(*Y_n,pred_(K)* − *Y_n,sim_*)^2^ using one of three objective functions: (i) the steady-state cluster size concentrations,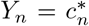 (ii) the cumulative steady-state cluster ter size distribution 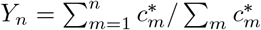; or (iii) the quasi-stationary cluster size distribution of fluctuating immobile domains, *Y_n_* = *A_n_*. The respective normalized errors are shown in Fig. 6A. We decided to use the value of *K* determined by the second method (cumulative size distribution) throughout this article because of its intermediate value and the relative sensitivity of the associated error to variations of *K*; note however that the differently determined values of *K* differed by at most 13% from that value and generally less than 10%.

**FIG. 6.**
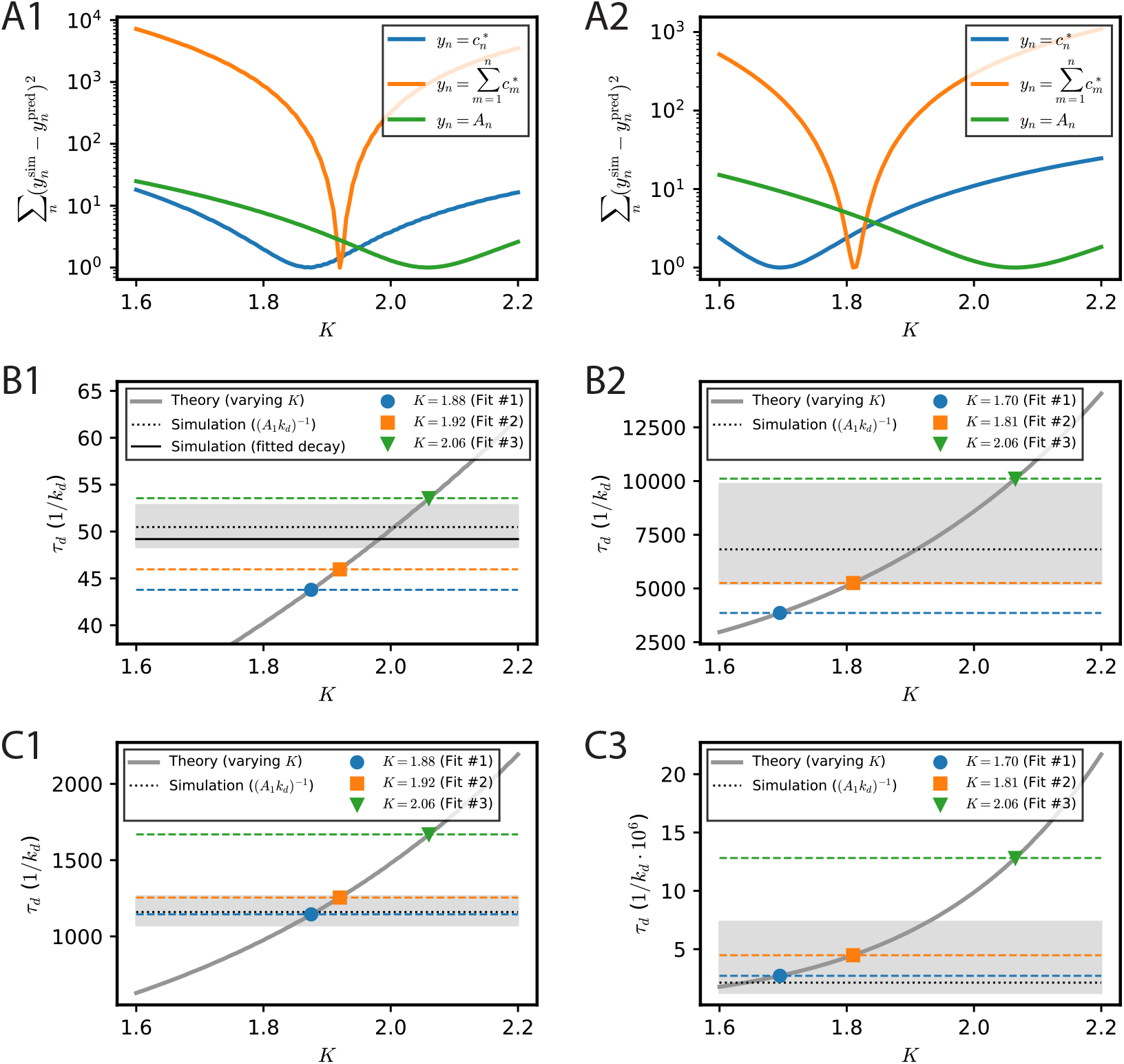
Determination and influence on predicted lifetimes of the Smoluchowski kinetic coefficient *K*. Left panels correspond to case I parameters, right panels to case II parameters. The optimal parameter *K* in the Smoluchowski equations is determined by best fitting the results of full numerical equations. (A) Sum of least-squared errors for the three objective functions indicated on the graph, as a function of the kinetic coefficient *K* in the mean-field description. Fitted values of *K* correspond to the respective minima of the errors. (B,C) Predicted cluster lifetime *τ_d_* based on the Smoluchowski rate equations as a function of *K* (thick solid grey line) and the value of *τ_d_* for the three optimal values of *K* determined according to the fits shown in panel A, for immobile (B) and diffusing (C) domains. The expected true cluster lifetime can be inferred from the value *A*_1_ of the quasi-stationary size distribution observed in the simulations and is shown (dashed black line) together with the 95% confidence interval obtained from bootstrap resampling, see Appendix C for details. In the case of fast particle turnover and for immobile domains (panel B1), the cluster lifetime can be measured directly from the particle-based simulations (thin solid black line).

The predicted cluster lifetimes as a function of *K* (thick solid grey line) are shown in Fig. 6B,C for immobile and diffusing domains, respectively. Note how small variations in *K* can give rise to considerably larger variations in the predicted cluster lifetime *τ_d_*. The expected cluster lifetime as inferred from the quasi-stationary size distribution observed in simulations (dashed black line) is superposed on all four panels, with the 95% confidence interval determined from bootstrap resampling shown in grey. Within this confidence interval, the expected cluster lifetime is consistent with the range of cluster lifetimes predicted from the different values of *K* determined using the three fit objective functions. Note that in the case of fast particle turnover and for immobile domains, Fig. 6B1, the explicitly measured cluster lifetime (thin solid black line) matches the expected cluster lifetime within the limit of precision of the latter.

